# Decoding Cell-type Specific Gene Regulation with CellReg3D by Integrating DNA Large Language Models and 3D Genome Structure

**DOI:** 10.64898/2026.11.01.763421

**Authors:** Xiaoyu Wang, Yumeng Zhang, Sihan Chen, Seiya Imoto, Geoffrey I. Webb, Athanasios V. Vasilakos, Fuyi Li, Kai Wang, Jiangning Song

**Affiliations:** Monash Biomedicine Discovery Institute and Department of Biochemistry and Molecular Biology, Monash University, Melbourne, VIC 3800, Australia; Monash AI Institute, Monash University, Melbourne, VIC 3800, Australia; Genome Center, Institute of Medical Science, The University of Tokyo, Minato-ku, Tokyo 108-8639, Japan; Collaborative Research Institute for Innovative Microbiology, The University of Tokyo, Bunkyo-ku, Tokyo 113-8657, Japan; Department of Data Science and Artificial Intelligence, Monash University, VIC 3800, Australia; Center for AI Research (CAIR), University of Agder (UiA), Grimstad, Norway; South Australian Immunogenomics Cancer Institute (SAiGENCI), Adelaide University, Adelaide, South Australia 5005, Australia; Department of Respiratory and Critical Care Medicine, the Fourth Affiliated Hospital of School of Medicine, and International School of Medicine, International Institutes of Medicine, Zhejiang University, Yiwu, 322000, China; Zhejiang Key Laboratory of Precision Diagnosis and Treatment for Lung Cancer, Yiwu, 322000, China; Zhejiang-Sweden Joint Laboratory on Tumor Immunology, Yiwu, 322000, China; Department of Veterinary Biosciences, Faculty of Science, The University of Melbourne, Parkville, Victoria, Australia

**Author notes:** Corresponding authors. Jiangning Song,; Kai Wang,; Fuyi Li,.

**Keywords:** Deep learning, gene expression, 3D genome, DNA language model, non-coding variant, long-range genomic interaction

## Abstract

Gene regulation is a core biological mechanism crucial for organismal growth and development. Recent advances in deep learning, coupled with increasing computational resources, have enabled large-scale modelling of gene regulatory processes. While numerous models have been proposed, effectively capturing long-range genomic interactions remains a central challenge. Here, we introduce CellReg3D, a multiscale and multi-modal deep learning framework designed to model cell-type-specific gene regulation by integrating local sequence patterns with long-range genomic interactions. Existing approaches often overlook the contribution of 3D genome architecture and face limitations in combining sequence-based features with chromatin interaction data. Specifically, CellReg3D overcomes these gaps by leveraging a DNA language model to encode sequence-level information and incorporating Hi-C-derived 3D chromatin interactions to capture chromatin spatial organization. We demonstrate that CellReg3D effectively learns epigenetic patterns and chromatin activity states, achieving state-of-the-art performance in predicting epigenetic marks and gene expression profiles across multiple cell types. Using interpretable methods, we illustrate how the model identifies cell-type-specific regulatory mechanisms and prioritizes functional non-coding variants. Furthermore, we apply CellReg3D to enhance the interpretation of expression quantitative trait loci (eQTLs) and uncover novel potential gene regulatory interactions. Overall, CellReg3D offers a integrative framework for decoding regulatory genomics, unifying sequence and 3D genome structure data to advance variant prioritization, disease mechanism discovery, and personalized functional genomics.

## Introduction

Gene regulation is a highly intricate process governed by interactions across multiple regulatory layers and diverse regulatory elements. Understanding its mechanisms is a fundamental question in molecular biology, given its central role in cellular differentiation, tissue development, and environmental response. Dysregulation of this process is often closely associated with the onset and progression of numerous diseases. As biological datasets continue to grow in both scale and complexity, a critical challenge lies in effectively integrating multimodal information to decipher gene regulatory mechanisms. Over the past decade, deep learning methods have revolutionized our ability to analyze gene regulation, becoming an indispensable tool in modern genomics research [1]. Gene expression is regulated at multiple levels, encompassing local epigenetic mechanisms—such as chromatin accessibility and histone modifications—and long-range chromatin interactions. It involves complex interplay between transcription factors and cis-regulatory elements like promoters, enhancers, and silencers, along with epigenetic modifications including DNA methylation, histone modifications, and chromatin remodeling, which collectively modulate gene accessibility [2]. Advances in diverse sequencing technologies, coupled with enhanced computational power, now enable deep learning approaches to analyze large-scale, multimodal biological datasets effectively.

Moreover, approximately 98% of the human genome is non-coding, and correspondingly, over 90% of single-nucleotide polymorphisms (SNPs) are located within these regions—areas primarily involved in epigenetic and gene regulatory processes [3]. Since the functional effects of such variants cannot be inferred from protein structure and experimental validation remains costly and lab-intensive, computational prediction has become indispensable. A widely used strategy involves simulating gene regulation by introducing *in silico* mutations and assessing their predicted functional impact. The accuracy of this approach hinges on precisely modelling gene regulatory mechanisms. Rapid advances in deep learning have significantly accelerated progress in variant effect prediction, drawing substantial research interest across genomics [1]. In recent years, a variety of deep learning models, including both supervised and unsupervised, have been applied to decipher complex gene regulatory patterns. Models trained on epigenetic and transcriptomic data, such as Borzoi [4], Enformer [5], Graphreg [6], Basenji2 [7], and Basenji [8], have markedly improved the prediction of genome function and variant effects. AlphaGenome [9], a recent model, further advances our understanding of genome function. In parallel, sequence-to-function models trained on long sequences and genetic tracks, including DeepSEA [10], Expecto [11], Sei [12], and Puffin [13], have been specifically designed to predict local functional elements and have demonstrated strong performance. Notably, GET [14], a foundation model tailored to human cell types, has emerged as a powerful tool for predicting regulatory elements.

Furthermore, numerous DNA large language models (LLMs), such as Caduceus [15], Nucleotide Transformer [16], DNABERT [17], and GROVER [18], have emerged as powerful tools for deciphering genomic grammar, providing deeper insights into the regulatory code of non-coding sequences. Another notable approach, Evo2 [19], is a large-scale foundation model trained via a two-phase strategy to capture genomic representations across both prokaryotic and eukaryotic organisms.

However, a significant limitation of existing models is their training on the reference genome alone, which may restrict generalizability to diverse genomic backgrounds. A pivotal challenge stems from the enormous combinatorial complexity of DNA; current technologies cannot exhaustively capture all possible sequence variations, leading to inherent data sparsity. While emerging DNA language models hold potential to address these constraints through their generative and predictive capabilities and show promise for genomic tasks, such as regulatory element prediction, they remain essentially black boxes for deciphering mechanistic gene regulatory processes.

In this work, we introduce CellReg3D, an interpretable, multiscale, and multimodal deep learning framework designed to predict epigenetic and gene expression profiles, as well as the effects of non-coding variants. While our primary goal is to build cell-type-specific predictors, conventional large-scale training paradigms often dilute cellular specificity by capturing shared rather than unique cell-type-specific features. Kathail et al. mentioned that multitask learning might allow the model to learn shared features rather than cell-type-specific features [20]. Therefore, developing a more interpretable model of cell-type specificity is essential to gain a more comprehensive understanding of gene regulation. Training is performed on a per-cell-line basis, enabling CellReg3D to learn cell-type-specific regulatory patterns. Furthermore, given that enhancer activity is known to depend on genomic distance [21], we incorporate 3D genome architecture into our model. This allows CellReg3D to explicitly account for the spatial interactions between regulatory elements, a critical factor often overlooked in sequence-only approaches.

To overcome the challenge of DNA data sparsity, our model combines a pre-trained DNA language model with cell-type-specific regulatory data to enable the inference of cellular states. The language model, trained on general DNA sequence representations, captures hierarchical patterns of regulatory tokens and functional motifs. These learned representations are then adapted within a cell-state-specific predictive architecture. Consequently, CellReg3D effectively captures epigenetic patterns and chromatin activity from DNA sequence and chromatin 3D structure inputs to predict epigenetic and gene expression profiles. This framework establishes a foundation for constructing biological digital twins, where individual cell-type models can integrate diverse cell-type-specific omics data. Its modular design also allows for the ready extension to new cell types, promising robust predictive performance across diverse biological contexts.

## Results

### Multiscale and multimodal design of CellReg3D

Our study comprises three major components: (i) the collection and processing of cell-specific datasets, (ii) the training of the deep learning model, and (iii) the downstream evaluation and analysis (**Fig. 1a)**. The dataset was constructed primarily using resources from the ENCODE database [22, 23] and the 4D Nucleome Project [24, 25]. Both follow standardized protocols with stringent quality control, ensuring high reliability and reproducibility of the data.

**Figure 1.**
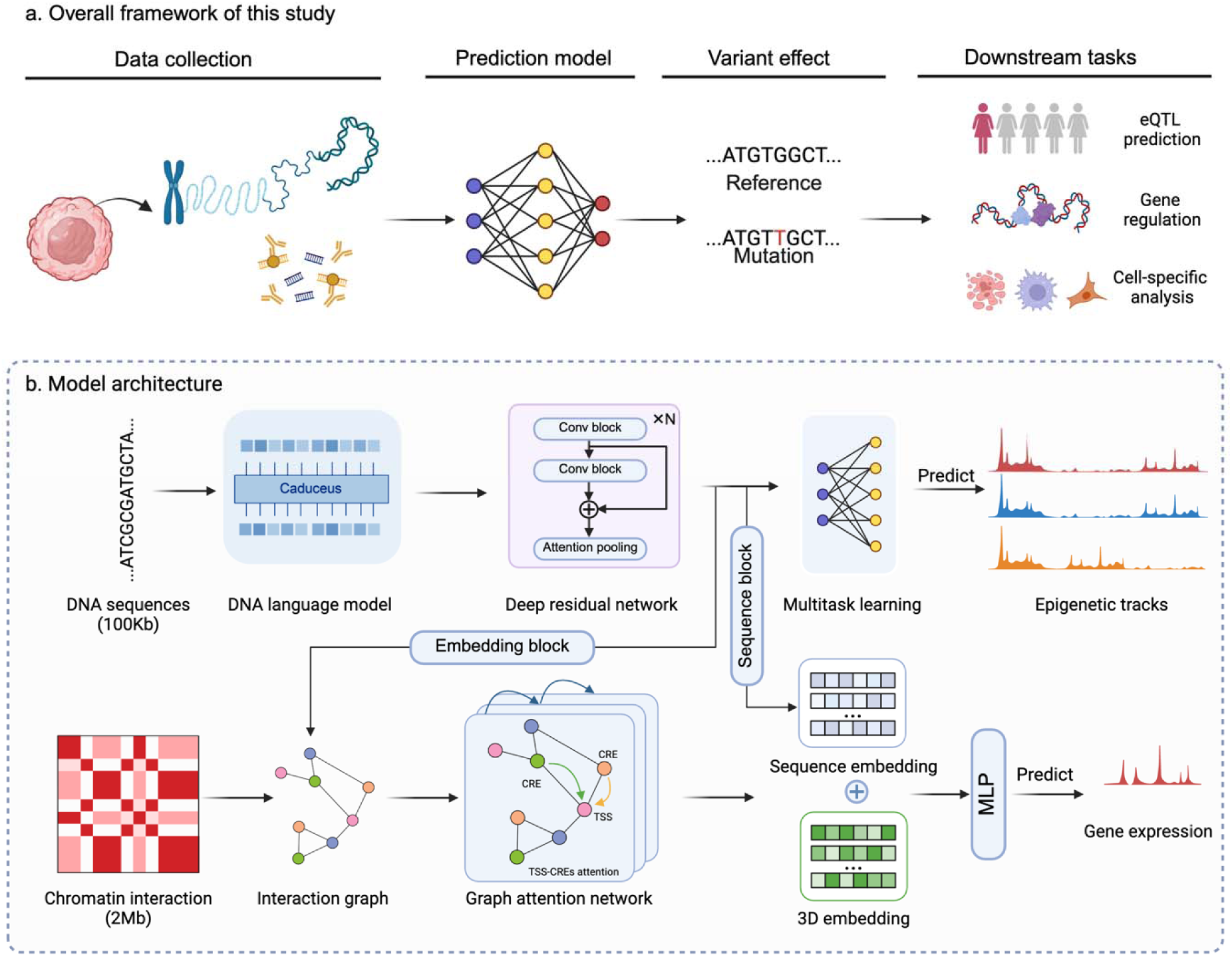
Framework of CellReg3D. a, Schematic of the CellReg3D pipeline for predicting genetic profiles and variant effects. The pipeline comprises four key components: data collection, predictive model construction, variant effect definition, and downstream analyses. b, model architecture of CellReg3D. DNA sequence and chromatin interaction information are used for the training process.

Based on these data, we developed a multiscale and multimodal deep learning model, termed CellReg3D, which integrates a large language model (LLM) with a graph attention network (GAT) [26] to predict epigenetic marks and gene expression from DNA sequences and 3D genome information (**Figure 1b; Extended Data Fig. 1;** see **Methods**). The model is designed to incorporate genomic information from regions up to 2 Mb away, substantially increasing its capacity to detect distal cis-regulatory elements.

### CellReg3D consists of three core components

i. an LLM-based Epigenetic Module: This module takes 100kb DNA sequences as input. To capture local epigenetic patterns, we employ a pre-trained DNA language model, Caduceus, which was trained on the human reference genome. Given the enormous combinatorial complexity of DNA sequences, such language models are essential for inferring cell-type-specific chromatin states. The sequence representations are further processed through a deep residual network to learn predictive local epigenetic patterns.
ii. 3D Genome Graph Attention Network (GAT) Module: This module explicitly models long-range spatial interactions between regulatory elements using chromatin conformation data (e.g., Hi-C).
iii. Integrated Epigenetic Prediction Module: The outputs from the sequence and 3D genome structure branches are integrated to predict a suite of multiple epigenetic profiles. The epigenetic prediction targets include key markers of chromatin state, such as chromatin accessibility (DNase-seq [27]) and multiple histone modifications (ChIP-seq [28]). Epigenetic modifications regulate gene expression by dynamically altering chromatin states. In this work, we modeled the chromatin states using eight epigenetic modifications, including DNase-seq, H3K4me3, H3K27ac, H3K36me3, H3K9me3, H3K4me1, H3K27me3, and H4K20me.

A key limitation of current sequence-based models is their underlying assumption of linear, distance-dependent regulatory efficacy. This contradicts experimental findings showing that gene regulation operates through nonlinear principles governed by 3D chromosomal organization, where mechanisms like chromatin looping can bypass linear genomic distance. Gene regulation occurs across multiple spatial scales: from local regulatory elements (*e.g.*, promoters, enhancers, and silencers), intermediate-scale epigenetic modifications (including DNA methylation and histone marks), through gene regulatory processes typically confined within topologically associating domains (TADs), up to higher-order architectures like chromatin compartments [29, 30]. To capture this multiscale nature, we implemented a deep neural network architecture capable of simulating the nonlinear dynamics of transcription. While most prior methods overlook 3D genome information, with GraphReg [6] being a notable exception for incorporating chromatin interaction information, our model integrates 3D genome information explicitly to complement sequence-based predictions. Specifically, we process 3D genomic features derived from Hi-C data within a 2 Mb region using a Graph Attention Network [26]. This module captures spatial interactions between cis-regulatory elements and transcription start sites. In parallel, DNA sequence information is processed and aggregated by the dedicated sequence modules. This dual-pathway design enables CellReg3D to jointly learn from both linear sequence context and nonlinear 3D genome structure relationships.

Our computational framework integrated eight well-characterized human cell lines spanning multiple tissue types and states: K562 (leukemia), GM12878 (lymphoblastoid), H1 and H9 (embryonic stem cells), HCT116 (colorectal cancer), HepG2 (liver carcinoma), HeLa-S3 (cervical adenocarcinoma), and IMR-90 (lung fibroblasts). These cell lines provide a comprehensive biological spectrum for analysis, encompassing normal, cancerous, and pluripotent stem cell types.

### *De novo* predictions of cell-type-specific epigenetic profiles and gene expression

We benchmarked CellReg3D against three leading methods for epigenetic profile prediction, including Enformer [5], GraphReg [6], and Basenji2 [7]. For a fair comparison, we adapted these models to accept the same inputs to predict epigenetic tracks. Predictions were evaluated across multiple cell lines using the Pearson correlation coefficient (PCC), the Spearman correlation coefficient (SCC), and the mean squared error (MSE) loss. Our model demonstrated statistically significant improvements over these benchmarks. Compared to Enformer, CellReg3D achieved mean Pearson correlation coefficient increases of 18.75% across the five cell types (two-sided Wilcoxon signed-rank test, *P* < 1.57×10^-10^; **Figure 2a and Methods**). We also observed significant gains over GraphReg (*P* < 1.14×10^-13^; **Figure 2b**) and Basenji2 (*P* < 3.66×10^-9^; **Figure 2c**). While Enformer and Basenji2 are designed for cross-cell-line modeling, their generalized architectures can lead to suboptimal performance on individual cell types. In contrast, CellReg3D can capture more precise, cell-type-specific regulatory patterns. Consistent superior performance is further demonstrated across multiple independent genomic datasets (**Extended Data Fig. 2, Supplementary file)**. We note that Borzoi [4] was also applied to our datasets; however, substantial differences in data scale and processing workflows precluded a direct and fair comparison with our method.

**Figure 2.**
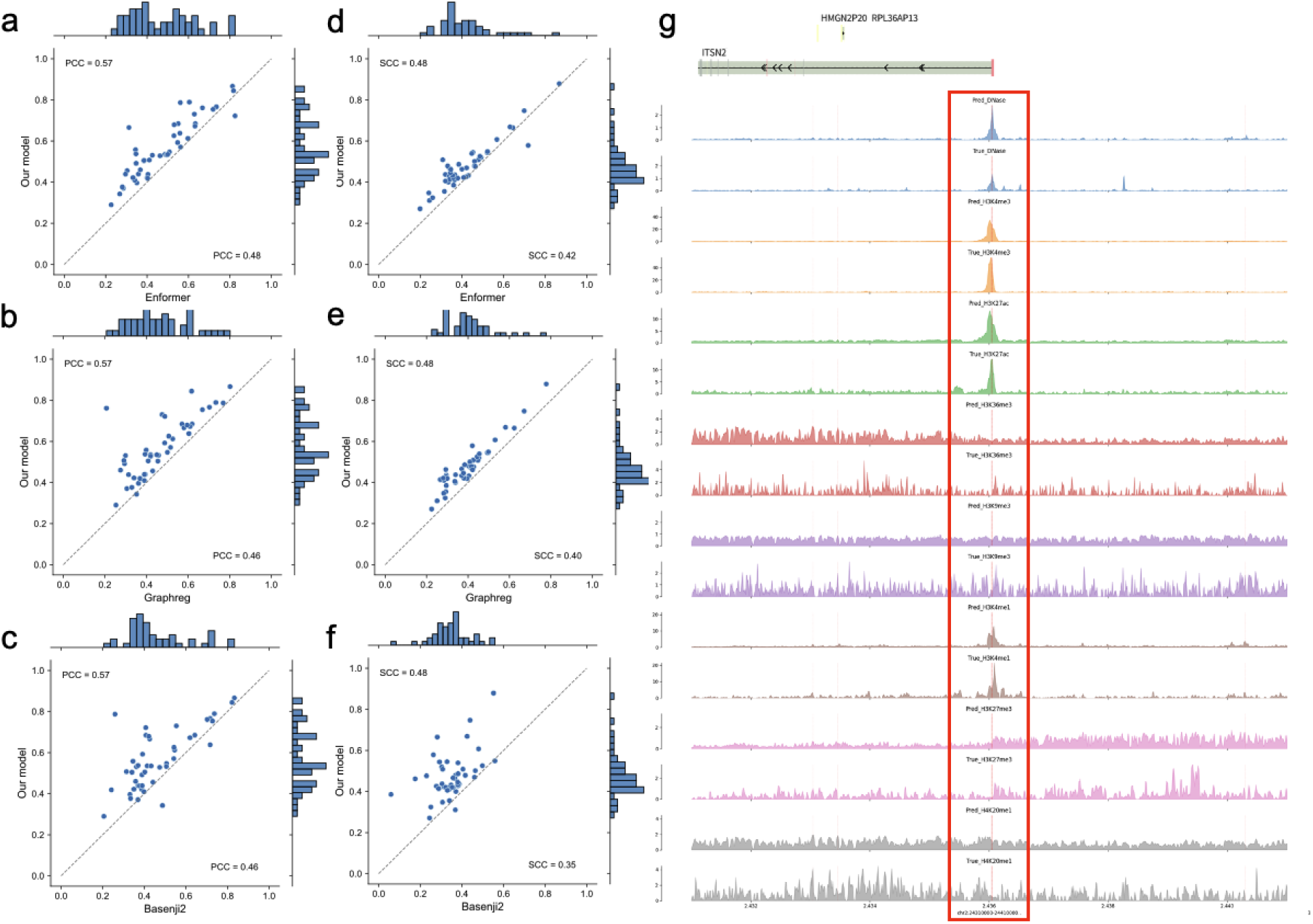
Performance comparison of CellReg3D. a-f, CellReg3D achieves higher PCC and SCC than Enformer, Graphreg, and Basenji2 in cell-type-specific evaluations. g, Representative example of predicted genetic tracks with ground truth (chr2: 24310000-24410000).

Our benchmarking was performed on a single-cell-line basis. Consequently, the performance metrics are not directly comparable to those reported in the original publications (*e.g.*, Enformer and Basenji2), which were evaluated across broader, multi-tissue cell line panels. CellReg3D outperformed these state-of-the-art methods based on the same datasets. Moreover, when evaluated using the Spearman correlation coefficient (SCC), CellReg3D achieved significant improvements over the competing models (**Figures 2d-f**). Superior performance was further validated using the MSE loss metric (**Extended Data Fig. 3**).

A qualitative assessment of predicted epigenetic tracks reveals close concordance with experimental ground-truth data, demonstrating CellReg3D’s ability to accurately recapitulate chromatin activity and epigenetic modification patterns (**Figure 2g, Extended Data Fig. 4**). The model’s robust quantitative performance is also summarized in dot-plot visualizations (**Extended Data Fig. 5-6)**. It is important to note that all reported results are from models trained on individual cell lines. Given the cell-type-specific nature of regulatory programs, performance may vary when models are trained on or applied to multi-cell-line datasets.

### Model ablation study

The performance improvements of CellReg3D are attributable to multiple factors. To systematically evaluate their individual contributions, we conducted a comprehensive ablation study focusing on four key components: (1) the 3D Genome GAT Module (Hi-C data), (2) the DNA language model, (3) the sequence-based module for gene expression, and (4) the multitask learning strategy for epigenetic patterns.

First, we evaluated the model’s performance with and without the 3D genome GAT module (Figure 3a). The results demonstrate that incorporating 3D genome information consistently improves performance. Second, we assessed the impact of the DNA language model by comparing three model variants: one trained using pre-trained LLM embeddings, one with a fine-tuned LLM, and a base model without any LLM component. The variant model with the fine-tuned LLM achieved significantly better performance (*P*-value < 0.05), indicating that adaptive sequence feature extraction is critical for optimal prediction (**Figures 3b-c**). This analysis revealed that the fine-tuned DNA language model contributed significantly to the model’s performance. Third, we evaluated the contribution of the sequence module. This component, which integrates the 1D regulatory context, consistently improved model performance, confirming its importance in capturing local regulatory grammar (**Figure 3d and h**).

**Figure 3.**
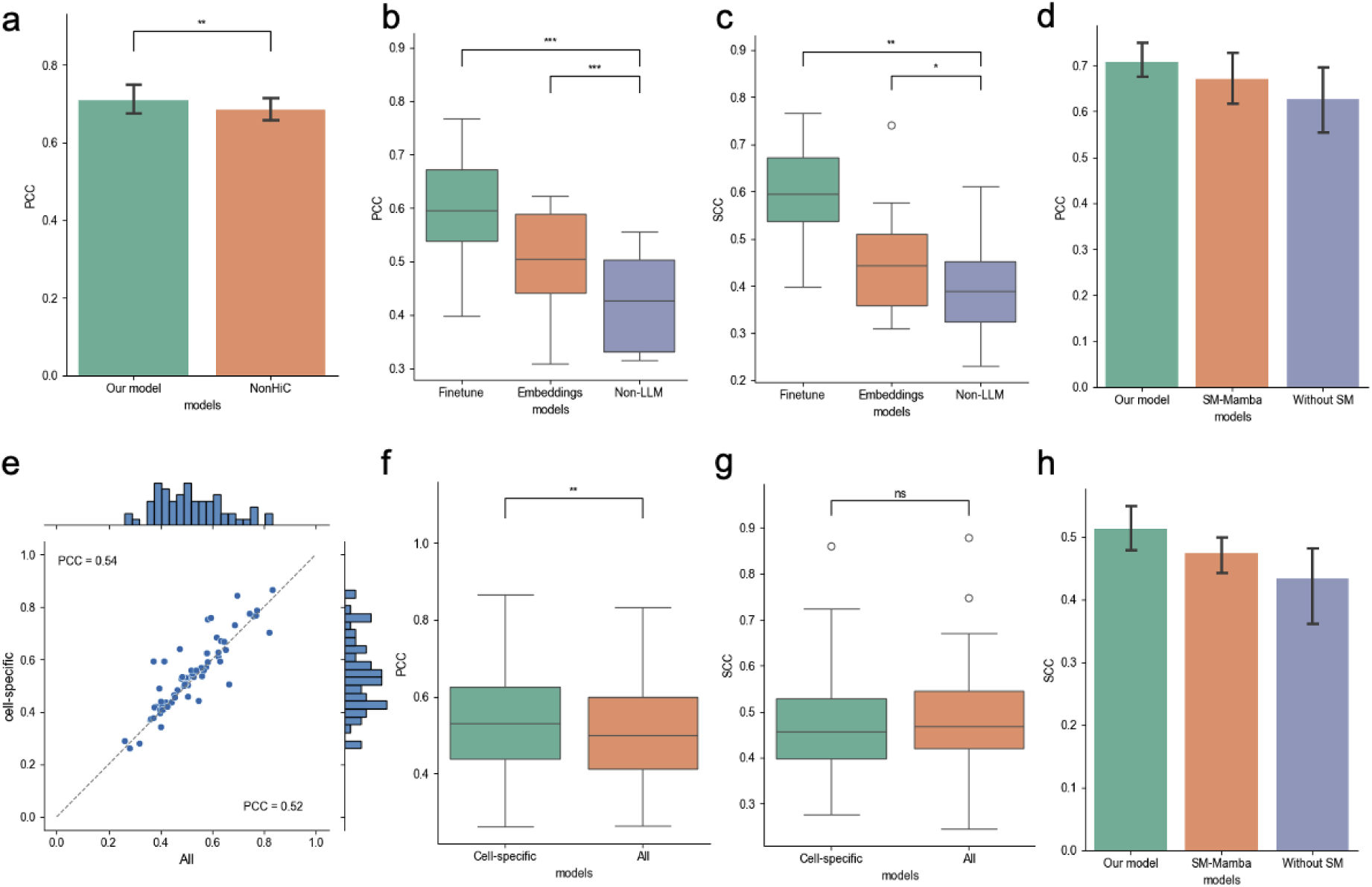
Model ablation study. a, Performance comparison of model variants: with Hi-C data and without Hi-C data. b and c, Performance comparison of model variants: without LLM, with LLM embeddings, and with the language model, evaluated using PCC and SCC metrics. * indicates statistical significance (p < 0.05), ** indicates statistical significance (p < 0.01), and *** indicates statistical significance (p < 0.001). d and h, Performance comparison of model variants with no sequence module, Mamba, or Transformer sequence modules based on PCC and SCC. e, Dot plot showing the performance of cell-specific models and the model trained on all cell lines. f and g, PCC and SCC results for cell-specific models and the model trained on all cell lines.

Moreover, we examined the impact of the multitask learning strategy by comparing our primary cell-type-specific model to the variant model trained on multiple cell types. In our framework, this approach did not yield significant performance improvements over the dedicated per-cell-line models on SCC, and it achieved lower performance on PCC (**Figures 3f-g**). Collectively, the ablation study demonstrates that the fine-tuned DNA language model and the 1D sequence module are essential for accurate gene expression prediction, whereas a multitask approach across cell types provided limited additional benefit within our architecture.

### CellReg3D recapitulates epigenetic modifications and chromatin activity

We systematically evaluated our model’s ability to recapitulate the epigenetic landscapes surrounding transcription start sites. Comparative visualization of predicted and experimentally derived chromatin feature tracks demonstrates that the model accurately captures spatial patterns of epigenetic modifications (**Figure 4**). Genomic regions with similar chromatin activity profiles are co-clustered based on their predicted epigenetic signatures, indicating that the model’s learned representations effectively encode biologically relevant spatial relationships.

**Figure 4.**
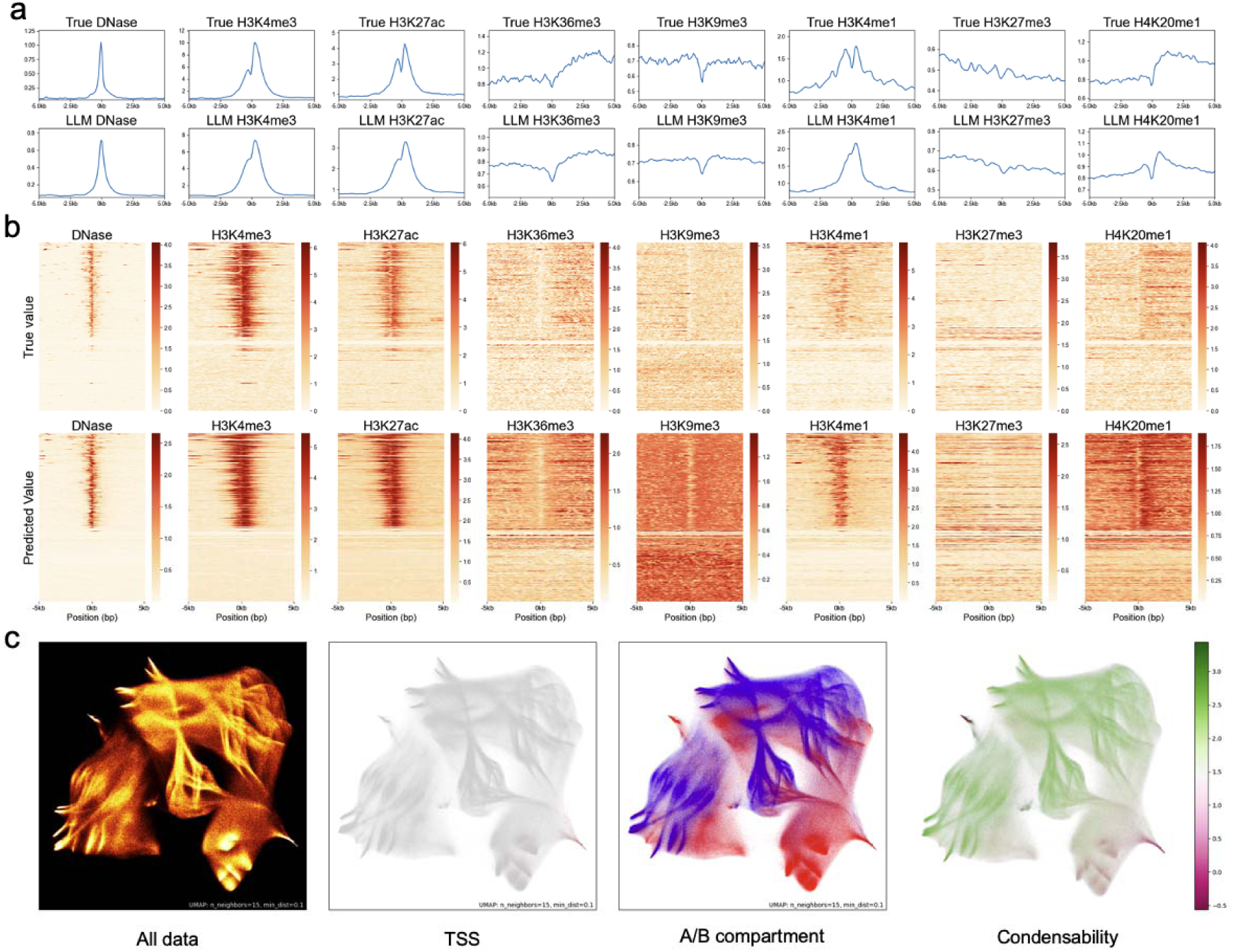
**a**. Average genetic track predicted by our model with the ground truth genetic track of chromosome 2 in TSS-proximal regions. **b**. Heatmap showing the genetic track of the ground truth (top) and model prediction (bottom). Color intensity corresponds to signal strength. **c**, UMAP projection of sequence embedding vectors based on the GM12878 cell, where each mark represents a 100bp DNA sequence.

To validate that the model captures meaningful epigenetic patterns, we analyzed average signal profiles for predicted versus experimental epigenetic tracks in TSS-proximal regions (±5 kb). Using aggregate line plots and sorted heatmaps for chromosome 2, we directly compared the spatial patterns of these signal profiles, confirming high concordance between predicted and ground-truth epigenetic data (**Figure 4a-b**).

To further investigate chromatin activity patterns, we visualized the hidden-layer embeddings for chromosomes 2 and 12 (test set) generated by our epigenetic module using UMAP (n_neighbors=15, min_dist=0.1). We then mapped three distinct functional genomic features onto the resulting embeddings: transcription start sites (from CAGE sequencing data), A/B compartments (derived from Hi-C data), and condensability scores (from Condense-seq [31]) for the GM12878 (**Figure 4c**) and H1 cell lines (**Extended Data Figure 7)**. Condense-seq quantifies the condensability of mononucleosomes, a property shown to correlate strongly with gene expression [31].

The UMAP projection of DNA sequence embeddings revealed clear spatial segregation of functional genomic elements: TSS regions clustered within high-activity domains, while A/B compartments indicative of active and inactive chromatin states formed distinct partitions. To quantitatively assess the biological relevance of these embeddings, we trained an XGBoost model [32] to predict condensability scores directly from the generated embeddings for chromosomes 2 and 12. Evaluated via 10-fold cross-validation, the XGBoost model achieved an average Pearson correlation coefficient of 0.61 and an average Spearman correlation coefficient of 0.59, outperforming both CNN and CNN–transformer hybrid baselines (**Table S4 and Extended Data Figures 8-9**). Notably, using only DNA sequence–derived features, the model achieved performance comparable to that of a prior method trained on multiple histone modification and DNA methylation profiles [31].

This integrative analysis systematically validates the biologically meaningful structure of the model’s latent space and enables a direct comparison with established genomic architectures.

### CellReg3D reveals cell-type-specific epigenetic and gene expression patterns

To evaluate the cell-type specificity of our model, we conducted a comprehensive cross-cell-type analysis, assessing both epigenetic mark prediction accuracy and gene expression modeling capabilities (**Figure 5a**). In this analysis, a model trained on one cell type was used to predict epigenetic states and gene expression in all other cell types, and performance was quantified by the average PCC for each genetic profile across different cell types in the regression task. The results demonstrate a strong self-prediction bias: all of the eight cell-line-specific models achieved their highest predictive accuracy when applied to predict their own cell type. This finding indicates that while CellReg3D captures fundamental, shared regulatory principles, it also retains a high degree of sensitivity to lineage-specific regulatory grammars. Detailed benchmarking performance metrics from this cross-cell-type evaluation are available in **Supplementary Table 5**.

**Figure 5.**
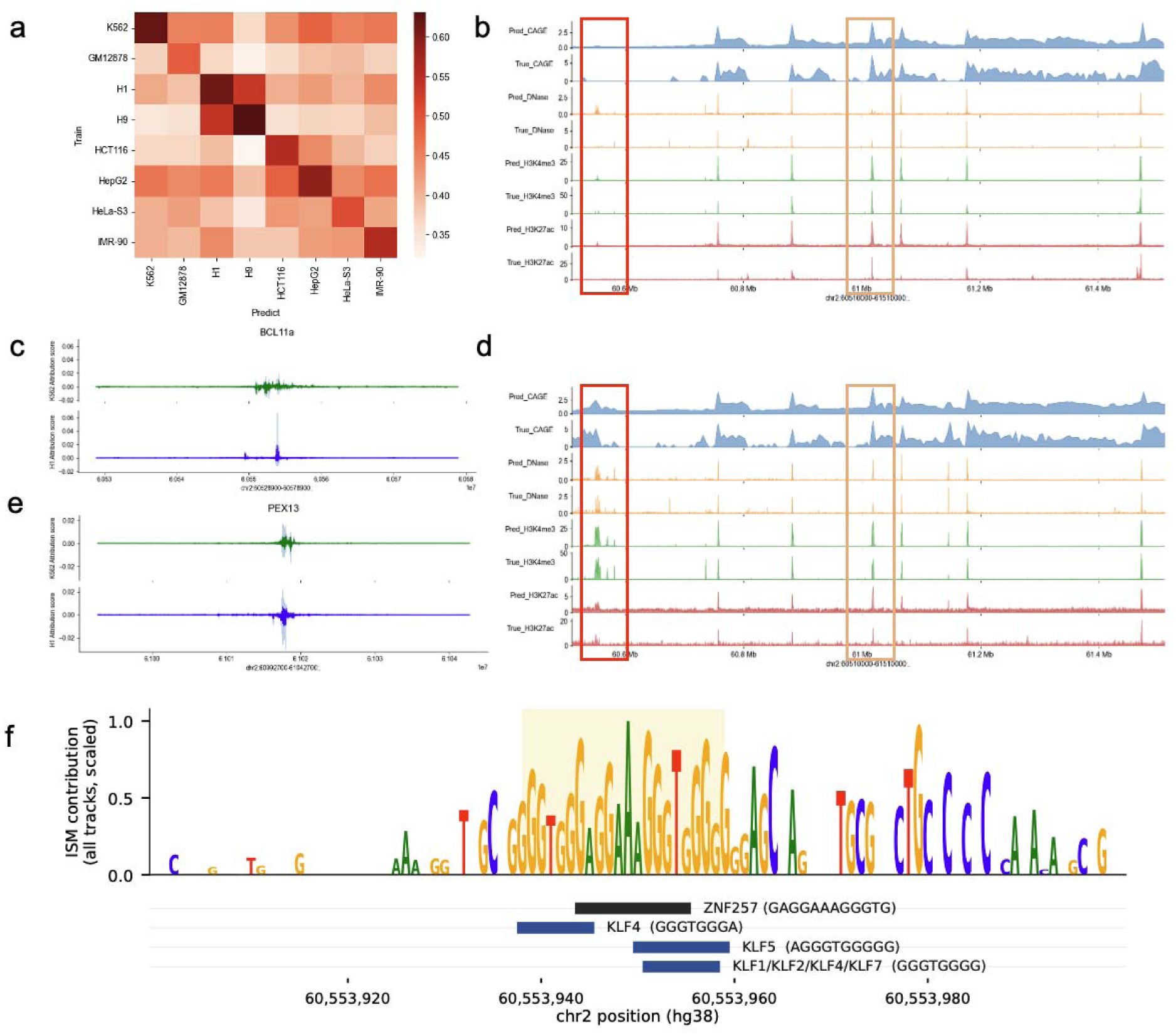
Cross-cell type-specific analysis. a, The performance evaluation of cross-cell type prediction is measured by the average PCC between predicted and target epigenetic and gene expression profiles. b, and d, predicted epigenetic and gene expression profiles and ground-truth values. The red box highlights the *BCL11A* gene, while the orange box indicates the *PEX13* gene. c and e, the contribution scores of *BCL11A* and PEX13 gene in K562 and H1 cell lines, respectively. The contribution scores were calculated using the input*×* gradient. f, ISM contribution scores and motif annotation across the BCL11A proximal promoter (chr2:60,553,900–60,554,000, hg38) in the H1 model.

To further investigate cell-type-specific regulatory mechanisms, we integrated multi-omic data, including Hi-C, gene expression, DNA accessibility, and key epigenetic marks (H3K4me3 and H3K27ac) for the K562 and H1 cell lines (**Figure 5d and e**). Notably, our model accurately recapitulated cell-specific gene expression patterns. For instance, the B cell lymphoma/leukemia 11A (*BCL11A)* gene, a key regulator of the fetal-to-adult hemoglobin switch in erythroid cells [33], was differentially expressed, showing higher expression in H1 than in K562. Consistent with this, the corresponding DNase-seq signals and activating epigenetic marks were lower in K562, suggesting a primary difference at the level of epigenetic regulation.

To test this hypothesis, we compared the contribution scores of a differentially expressed gene, *BCL11A*, with those of *PEX13* (a gene with similar, conserved expression levels across cell types) (**Figures 5b and c**). *PEX13* encodes an essential mammalian peroxin for the import of proteins containing a peroxisome targeting signal 1, a broadly conserved cellular mechanism across multiple cell types. The analysis revealed distinct regulatory patterns: *BCL11A* exhibited more focused, cell-type-specific contribution scores in the H1 cell line compared to the K562 cell line, whereas the distribution of the contribution scores for *PEX13* was highly similar between the two cell types.

To investigate the mechanisms of cell-type-specific gene expression, we used *BCL11A* as a case study, a gene exhibiting distinct expression patterns between H1 embryonic stem cells and K562 erythroleukemic cells. *BCL11A* is robustly expressed in H1 but markedly suppressed in K562, aligning with its established role as a transcriptional regulator in hematopoietic lineage commitment. Our model successfully captured this cell-type-specific differential expression, demonstrating its ability to identify key lineage-determining factors. Importantly, *BCL11A* is located on chromosome 2, which was held out from training for all cell-type-specific models; thus, none of the models had been exposed to the epigenomic or expression signals at this locus during training. To elucidate the regulatory logic underlying *BCL11A*’s expression, we applied an *in silico* saturation mutagenesis (ISM) score to quantify the nucleotide-resolution impact on predicted epigenetic profiles. In our analysis, the *BCL11A* locus showed a higher relative contribution score in the H1 model than in the K562 model. The region with the highest attribution was a GA-rich sequence (chr2:60,553,942– 60,553,965), which most closely matched motifs of several zinc-finger transcription factors, spanning the KLF and ZNF families. Given the distinct cellular contexts of H1 and K562, differences in transcription factor availability may contribute to cell-type-specific sequence patterns underlying chromatin state prediction. For example, KLF4, a reprogramming factor associated with pluripotency [34], has been shown to recruit the SWI/SNF chromatin-remodeling complex to increase chromatin accessibility [35]. This provides a potential mechanistic basis for cell-type-specific chromatin regulation.

### Functional insights from model interpretability

To elucidate our model’s decision-making process and identify key functional genomic elements, we performed comprehensive model interpretability analyses. These computational approaches enable the efficient prioritization of candidate functional elements and variants compared to purely experimental methods. Specifically, we calculated input gradients as contribution scores for each genomic region, yielding cell-type–specific importance maps for all target regions. We then validated the biological relevance of these scores through two downstream tasks: enhancer-gene pair identification and eQTL prioritization.

For enhancer-gene interaction prediction, we used validated enhancer candidates derived from CRISPRi-FlowFISH datasets [36], which experimentally perturb putative enhancers in K562 cells and quantify their effects on target gene expression. Candidate enhancers for each gene were ranked by their model-derived contribution scores, and performance was evaluated using the gene-wise area under the precision-recall curve (AUPRC). Across 27 target genes, our model-derived contribution score-based ranking outperformed the distance-to-TSS baseline (**Figures 6a-b**). While the Activity-by-Contact (ABC) score [36] achieved a higher performance on this task (**Extended Figure 10**), our method provides a more general and interpretable framework rather than a task-optimized heuristic for enhancer–gene linking. Importantly, our approach still achieved competitive performance for a subset of genes and offers complementary interpretability, enabling cross-validation of predicted enhancer-gene links and supporting other downstream analyses.

**Figure 6.**
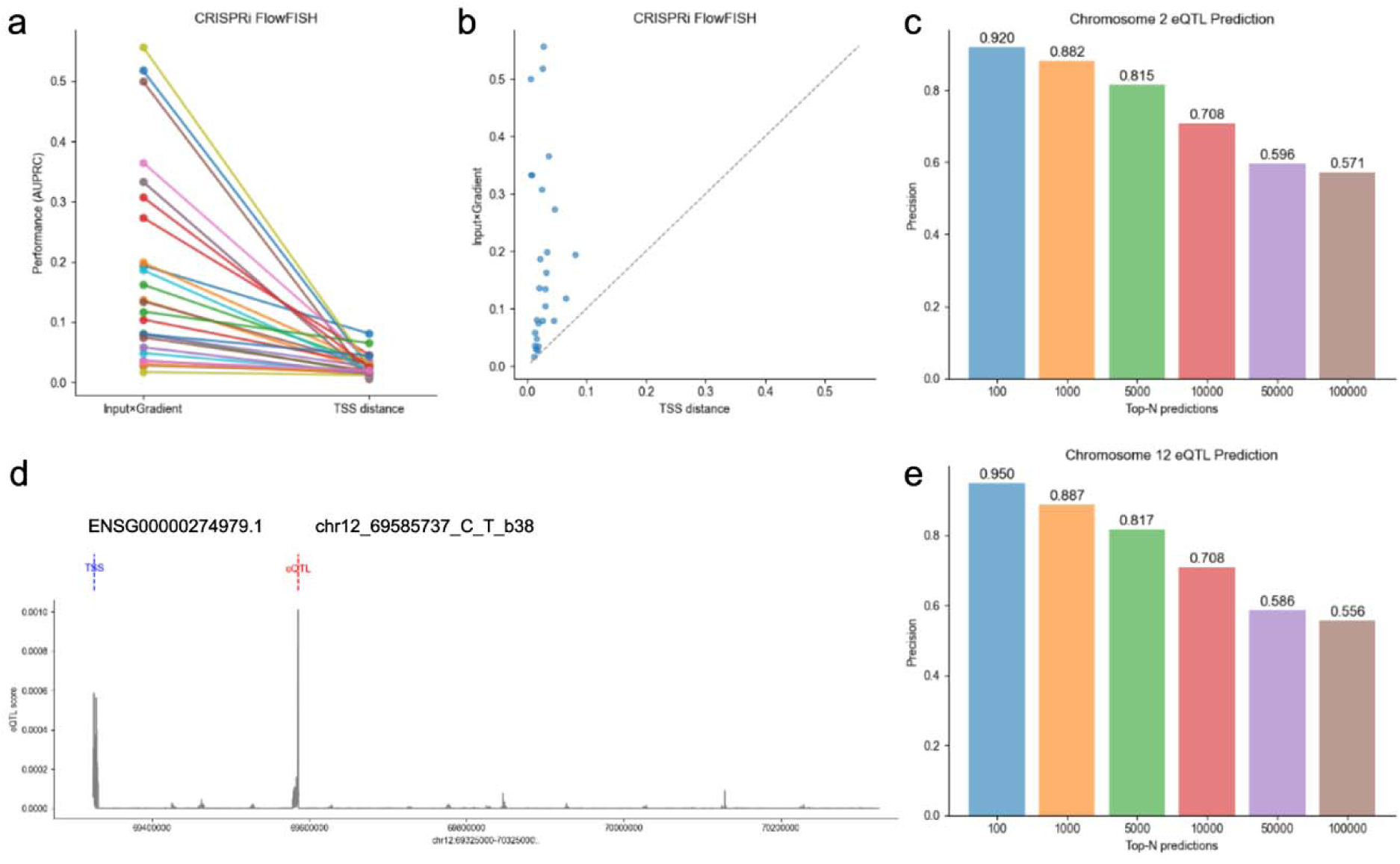
Downstream task evaluation. (a) Line plot comparing enhancer identification performance between contribution scores and a distance-based method. (b) Dot plot illustrating the AUPRC results for enhancer identification. (c, e) eQTL prediction performance on chromosomes 2 (c) and 12 (e). (d) An example showing contribution scores, the corresponding eQTL, and its target gene.

For eQTL prioritization, we constructed a dataset from GTEx whole-blood eQTLs [37] (positive variants) and randomly selected genetic variants from the 1000 Genomes Project [38] (negative controls). In this task, we focused on the precision of prioritized variants. As such, model performance was assessed by measuring the precision (the fraction of true eQTLs) among variants ranked highest by our model’s contribution scores. We calculated precision at multiple top-N thresholds (top 100, 1,000, 5,000, 10,000, 50,000, and 100,000 variants). The GM12878 model demonstrated strong enrichment for functional eQTLs, achieving precisions of approximately 0.82 among the top 5,000 ranked variants and 0.71 among the top 10,000 ranked variants (**Figures 6c and 6e**). This performance was consistent across the held-out test chromosomes 2 and 12, indicating robust generalization.

To illustrate the model’s capability to prioritize functional eQTLs, we examined a known whole-blood eQTL on chromosome 12 (linked to the gene ENSG00000274979.1). Our model assigned a high contribution score to the genomic region surrounding this variant, confirming that our interpretability-driven framework can successfully identify and prioritize validated regulatory variants.

## Discussion

In summary, we present CellReg3D, a multiscale, multimodal deep learning framework that predicts epigenetic and gene expression profiles to support the functional interpretation of noncoding genetic variation. Our model outperforms current state-of-the-art methods on focused datasets and successfully captures cell-type-specific epigenetic modification and gene expression patterns. It accurately identifies key genomic features, including transcription start sites and chromatin condensability, and employs explainable techniques to interpret the biological principles underlying differential gene expression. By analyzing feature contribution scores, CellReg3D can pinpoint candidate regulatory elements likely to drive expression differences.

These findings should be interpreted within the context of several methodological and practical limitations. While CellReg3D demonstrates robust performance on histone modification data, its training corpus could be expanded to incorporate additional regulatory layers, such as transcription factor binding and DNA methylation patterns. Furthermore, the resolution of our gene expression predictions is constrained by the inherent granularity of input 3D chromatin data (typically at the 1-5 kb scale for Hi-C versus 1 bp resolution for sequence data), thereby creating a fundamental bottleneck in spatial precision.

A primary challenge is the computational demand of processing large genomic inputs. While current models integrate sequences on the order of millions of nucleotides as input, human chromosomes exceed 100 million base pairs. Moreover, regulatory interactions can span vast chromatin distances. Accordingly, processing whole-genome sequences for accurate variant effect prediction remains a significant computational challenge.

Future developments of CellReg3D could integrate additional layers of biological specificity, such as cell-type- and tissue-specific regulatory information, including lineage-determining transcription factors and high-resolution spatial chromatin organization patterns, to improve the model’s generalizability across diverse tissues and conditions. Incorporating principles of nuclear organization [39] would further refine the modeling of higher-order chromatin architecture. Moreover, training on individual-level genomic and epigenomic data is a crucial next step for advancing precision medicine, as such personalized models could achieve greater disease specificity and enhanced clinical applicability and utility.

## Methods

### Model architecture

Our proposed CellReg3D architecture employs a dual-branch framework to integrate both 1D DNA sequence information and 3D structural genomic information. The first branch utilizes a large DNA language model to generate embeddings of DNA sequences, capturing long-range contextual patterns and latent regulatory semantics. The second branch processes 3D interaction graphs constructed from Hi-C interaction data, representing topological associations between genomic loci. For the first branch, we leverage a pre-trained large-scale DNA language model, named Caduceus, to encode 100k DNA sequences into context-aware embeddings. We implemented the Caduceus DNA language model using pretrained weights from Hugging Face’s Transformers library, which was originally trained on a reference genome using a masked language modeling approach. The model structure includes the BiMamba module to capture long-range interactions in DNA sequences.

Then, a deep residual network captures long-range interactions and reduces the feature dimensionality from 100 kb to 1,000 bins. Consequently, each bin in this representation corresponds to a 100 bp sequence segment. In the second branch, Hi-C contact matrices are converted into weighted graphs, where nodes represent 5kb genomic bins and edges denote interaction frequencies. The feature of each node is derived from the hidden layer of the LLM-based epigenetic model. The Graph Attention Network was adopted to capture the interaction between different chromatin regions. The interaction graph was defined as *G(V, E, A)*. Here, *V* is the node set, *E* is the edge set, and *A* is the adjacency matrix whose elements encode the corresponding edge weights. The attention coefficient *a_ij_* between the nodes i and j is computed as:

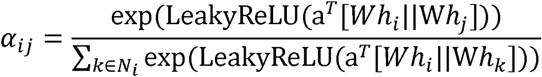

*h_i_* and *h_j_* are the input feature vectors of nodes *i* and j, respectively, and W is the learnable weight matrix. *N_i_* is the set of neighbor nodes (chromatin regions) connected to node *i* in the graph G.

The gene expression prediction task considers a 2 Mb genomic context surrounding each gene, allowing the model to account for long-range enhancer-promoter interactions. Additionally, to model the linear dependencies within the regulatory code, we also implemented a sequence model based on a transformer architecture. The detailed model structure is included in the Supplementary material.

### Input and output pre-processing for the model

The model requires two primary inputs: DNA sequence and Hi-C data. The genomic sequences were sourced from the human reference genome assembly GRCh38 (hg38) and subjected to trimming of low-confidence ends. The target genomic tracks include gene expression data profiled by CAGE sequencing and histone modification profiling by ChIP-Seq.

CAGE sequencing data in BAM format were obtained from the ENCODE and FANTOM5 consortium. Consistent with previous studies, to generate genome-wide coverage tracks, we converted BAM files to the bw format using the Basenji2 processing pipeline. Then the CAGE gene expression values were transferred as *log*(*x*+1). Bigwig files containing histone modification profiles were obtained from the ENCODE project. All genomic data in this study were aligned and analyzed based on the hg38 human reference genome. Genomic coordinates were converted from the hg19 to hg38 reference assembly using the pyliftover package, which was implemented based on the UCSC LiftOver algorithm.

To ensure rigorous evaluation of our model’s generalizability, we partitioned the genomic data into training, validation, and test sets using a chromosome-based split strategy. We partitioned the data by chromosome, using chr1 and chr10 for validation, chr2 and chr12 for testing, and the remaining autosomes for training. The reference genome was tiled into segments of 100 kb with a step size of 100K-bp, generating non-overlapping genomic regions for epigenetic profile prediction. We optimized the model parameters by minimizing the Poisson negative log-likelihood loss function, which has been widely used for genomic data. The loss function is defined as:

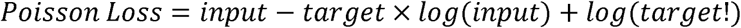

The model was developed based on PyTorch and the PyTorch Geometric (PyG) library, and training was performed on an NVIDIA Tesla A100 GPU, employing an early stopping strategy to mitigate overfitting. For epigenetic feature training, we used the AdamW optimizer with a learning rate of 5e-6 and a batch size of 2. For gene expression prediction, the batch size was reduced to 1, while all other optimizer settings remained consistent.

### Performance evaluation metrics

Model performance was comprehensively evaluated using three complementary metrics: Pearson correlation coefficient (PCC) to assess linear relationships between the ground truth and the predicted genomic tracks, Spearman correlation coefficient (SCC) to measure monotonic associations, and mean squared error (MSE) to quantify prediction accuracy. For downstream task evaluation, the area under the precision–recall curve (AUPRC) was used to assess model performance. The Pearson correlation coefficient is defined as:

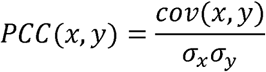

Where *x* and *y* represent the predicted and experimentally observed values, respectively, and *cov*(*x, y*) denotes their covariance, and σ*_x_* and σ*_y_* indicate the standard deviation of predicted genomic tracks and ground truth. This metric ranges from −1 to 1, with 1 indicating perfect linear agreement. Similarly, SCC evaluates monotonic relationships by computing PCC on rank-transformed data:

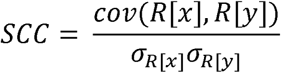

where *R[x]* and *R[y]* are the ranks of variables *x* and *y*, and *cov*(*R[x], R[y]*) is the covariance of the rank variables. σ*_R[x]_*and σ*_R[y]_* are the standard deviations of ranks.

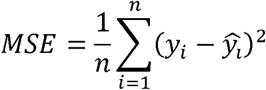

where *y_i_* denotes the true value of the i-th sample, *y_i_* is the predicted value, and n is the total number of samples. The MSE measures the average squared difference between the predicted and true values, with smaller values indicating better predictive performance.

### Model implementation

We compared our model with the three state-of-the-art methods for epigenetic and gene expression profiles prediction, including Enformer, Graphreg, and Basenji2. Early stopping was applied based on validation loss to prevent overfitting.

#### Enformer

The Enformer was implemented based on the PyTorch version (https://github.com/lucidrains/enformer-pytorch). It’s trained on our cell line-specific epigenetic and gene expression datasets. Enformer was originally trained on 196kb DNA sequences and 128bp resolution; we modified the Enformer model to support 100kb DNA sequences and 100bp resolution.

#### Graphreg

The GraphReg model was implemented in TensorFlow. To ensure a fair comparison, we maintained identical input configurations across GraphReg and CellReg3D, including data preprocessing steps and feature representations. The optimizer hyperparameters were kept identical to those used in the original model: an Adam optimizer with a learning rate of 2e-4 and a decay rate of 1e-6.

#### Basenji2

For benchmarking purposes, we also included Basenji2 in our performance comparison using its official implementation from GitHub (https://github.com/calico/basenji). The code is based on TensorFlow. The optimizer was configured with a learning rate of 2e-4 and a decay rate of 1e-6.

#### Borzoi

The Borzoi was implemented based on the TensorFlow version provided on the official GitHub (https://github.com/calico/borzoi). The parameters are taken from the Params.json file, where the input sequence length is 524,288 bp, and the resolution is 32 bp. The learning rate is 0.00001. The final output is generated solely by the human-specific heads of the model.

#### Model variations

For the model ablation study, we implemented several model variants based on the PyTorch framework. All ablation experiments were conducted by modifying specific components while keeping the remaining network architecture and training settings identical.

For the 3D genome module, two variants were developed to evaluate the contribution of 3D genome information: (1) a model with a Hi-C branch, which uses a GAT to integrate the Hi-C data, and (2) a variant without the Hi-C branch, where the GAT module was removed.

To evaluate the impact of the DNA language model, three variants were examined: (1) Fine-tuned LLM, in which all parameters of the pretrained DNA language model were fine-tuned; (2) embeddings, where the architecture remained the same but parameters were kept frozen; and (3) Non-LLM, which employed one-hot encoding for input sequences instead of learned embeddings.

For the sequence module (SM), three variants were constructed to evaluate its contribution: (1) SM-Transformer, which employs Transformer encoder layers configured with a hidden dimension of 64 and 8 self-attention heads; (2) SM-Mamba, implemented based on Mamba [40] and its code (https://github.com/state-spaces/mamba); and (3) a variant without the sequence module, in which this component was removed.

### Condensability prediction

To evaluate whether the model captures chromatin states, we constructed a condensability-prediction task using data from Park et al. [31]. To ensure data independence and fair evaluation without information leakage, the model was trained using data from chromosomes 2 and 12, which were held out as the test set. The genome was segmented into non-overlapping 10 kb bins.

CellReg3D-derived embeddings were used as input features, and the model was trained to predict condensability as a continuous variable, which serves as a quantitative measure of chromatin condensation for each genomic region. The model’s performance was assessed using the Pearson correlation coefficient (PCC) and the Spearman correlation coefficient (SCC) to quantify linear and rank-order agreement between predictions and true values. Model evaluation was performed using a repeated 10-fold cross-validation procedure, repeated 10 times, to obtain stable and unbiased performance. Multiple models, including XGBoost, CNN, and CNN+Transformer, were constructed for the condensability prediction task. Detailed descriptions of model architectures and implementation details are provided in the Supplementary Materials.

### Model interpretation

Interpreting language models remains a challenge. Two interpretation methods have been used in this study: *In silico* saturation mutagenesis (ISM) and Input × Gradient. The model interpretation is implemented using the Captum package [41]. Contribution scores were computed by integrating multiple epigenetic and gene expression tracks. The score for each target bin was defined as the average contribution across the target bin itself, along with two adjacent bins upstream and downstream.

### Enhancer prediction

We extracted 2-Mb DNA sequences centered at the target transcription start site to compute the Input×Gradient contribution scores. For each enhancer, we aggregated the contribution scores within a 4 kb genomic window centered on the enhancer midpoint, encompassing 2 kb upstream and 2 kb downstream. In addition, we calculated the relative distance between the midpoint of each enhancer and its corresponding target transcription start site for comparative analysis. For AUPRC calculation at the gene level, we excluded enhancers located more than 1 Mb from the target transcription start site. In addition, genes without experimentally validated enhancers and genes with fewer than 10 enhancer samples were removed to ensure reliable performance estimation.

### eQTL effect prediction

eQTLs were obtained from the GTEx portal, using the publicly available whole blood dataset. Negative samples were constructed by randomly selecting an equal number of variants from the 1000 Genomes Project on the same chromosomes. Data from two chromosomes were included in this analysis. Chromosome 2 contains 191,507 eQTLs and an equal number of negative variants, while chromosome 12 contains 124,643 eQTLs and an equal number of negative variants. The contribution score of each variant was calculated based on the GM12878 cell line model using the input × gradient method, measuring the contribution of each variant to the corresponding target gene.

### Epigenetic track visualization

To compare the true epigenetic track with the predicted track around the transcript start sites, we visualise the eight epigenetic modifications based on chromosome 2, which is in the independent test dataset. Transcription start sites were obtained from GENCODE, and the corresponding epigenomic tracks were visualized across a 10 kb region flanking each TSS, including 500 bp upstream and 500 bp downstream.

### Embedding visualization

The epigenetic module embeddings are obtained from the hidden layer of the model. We used UMAP [42] to visualize the distribution of embeddings of 100bp. TSS annotations were derived from GENCODE annotations and CAGE data, while A/B compartment information was obtained from ENCODE (K562: ENCFF699RSL; GM12878: ENCFF412CDH) and 4D data portal (H1: 4DNFIK34GR2J) Hi-C data. The chromosome-wide condensability of GM12878 and H1 cells was retrieved from the GEO database under accession number GSE252941 [31].

## Data availability

The Accession ID of the dataset can be found in the supplementary material. The blacklist of hg38 was downloaded from ENCODE (ENCFF356LFX.bed.gz). While the reference genome was downloaded from ENCODE.

## Code availability

The code for CellReg3D and the model weights are deposited on GitHub (https://github.com/yanwu20/CellReg3D).

## Acknowledgement

We acknowledge the ENCODE Consortium, the ENCODE production laboratories, and the 4D Nucleome (4DN) Consortium and participating laboratories for generating the datasets used in this study.

