## Supplementary files for "Decoding Cell-type Specific Gene Regulation with CellReg3D by Integrating DNA Large Language Models and 3D Genome Structure"

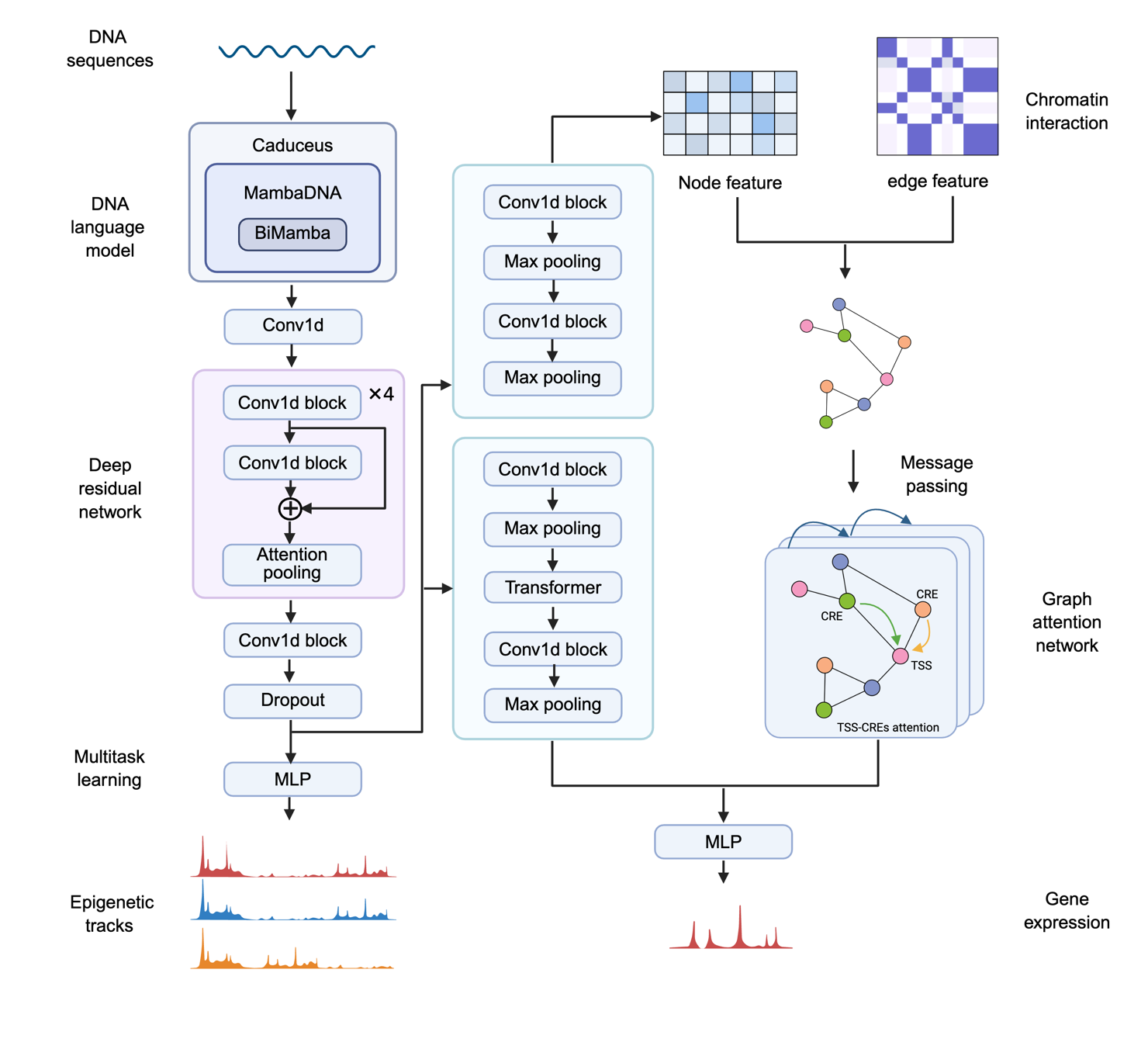

Figure S1. The framework of our model.

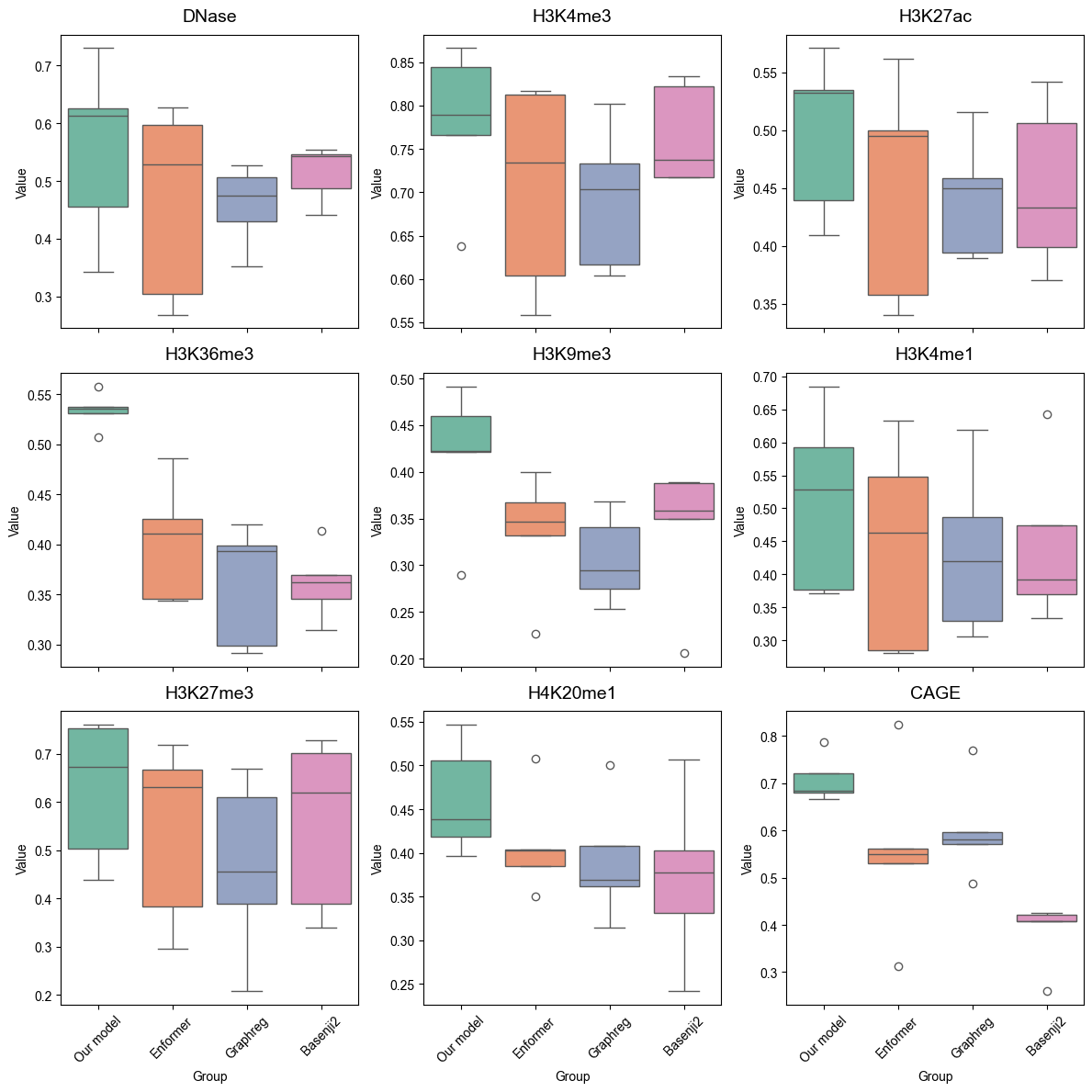

Figure S2. Performance comparison between our model and other state-of-the-art methods based on Pearson correlation coefficients for different genomic profiles.

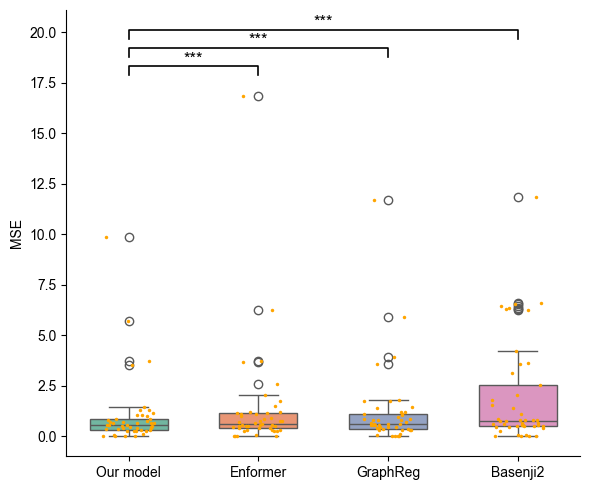

Figure S3. Distribution of mean squared error (MSE) values for models evaluated on five cell lines. *** indicates statistical significance (*p* < 0.001).

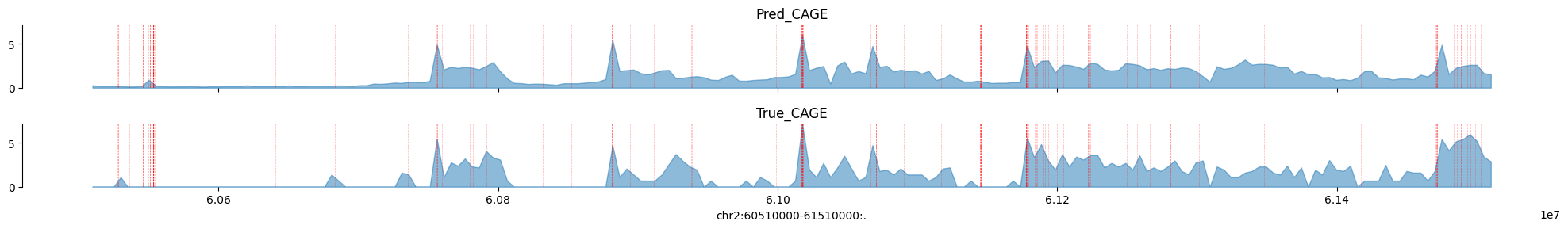

Figure S4. Representative example of predicted and ground-truth CAGE profiles. The red line indicates the transcription start sites.

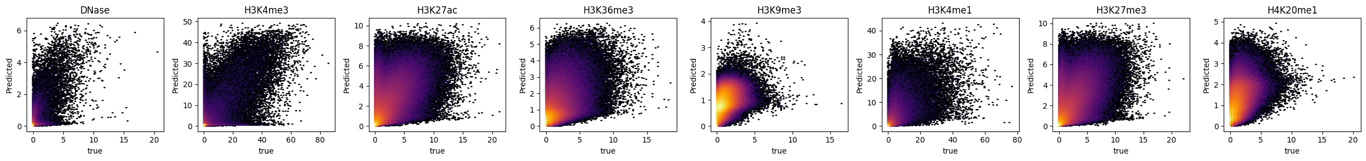

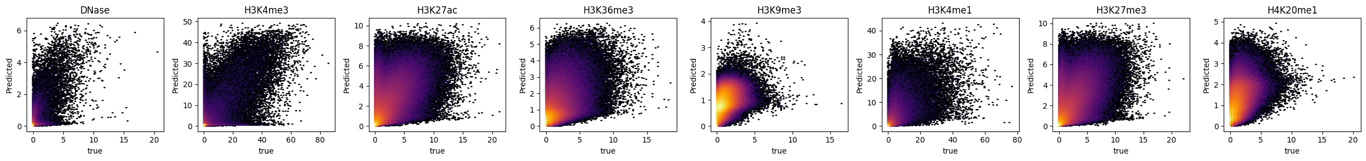

Figure S5. Comparison between predicted and experimentally measured DNase-seq and ChIP-seq coverage in the K562 cell line.

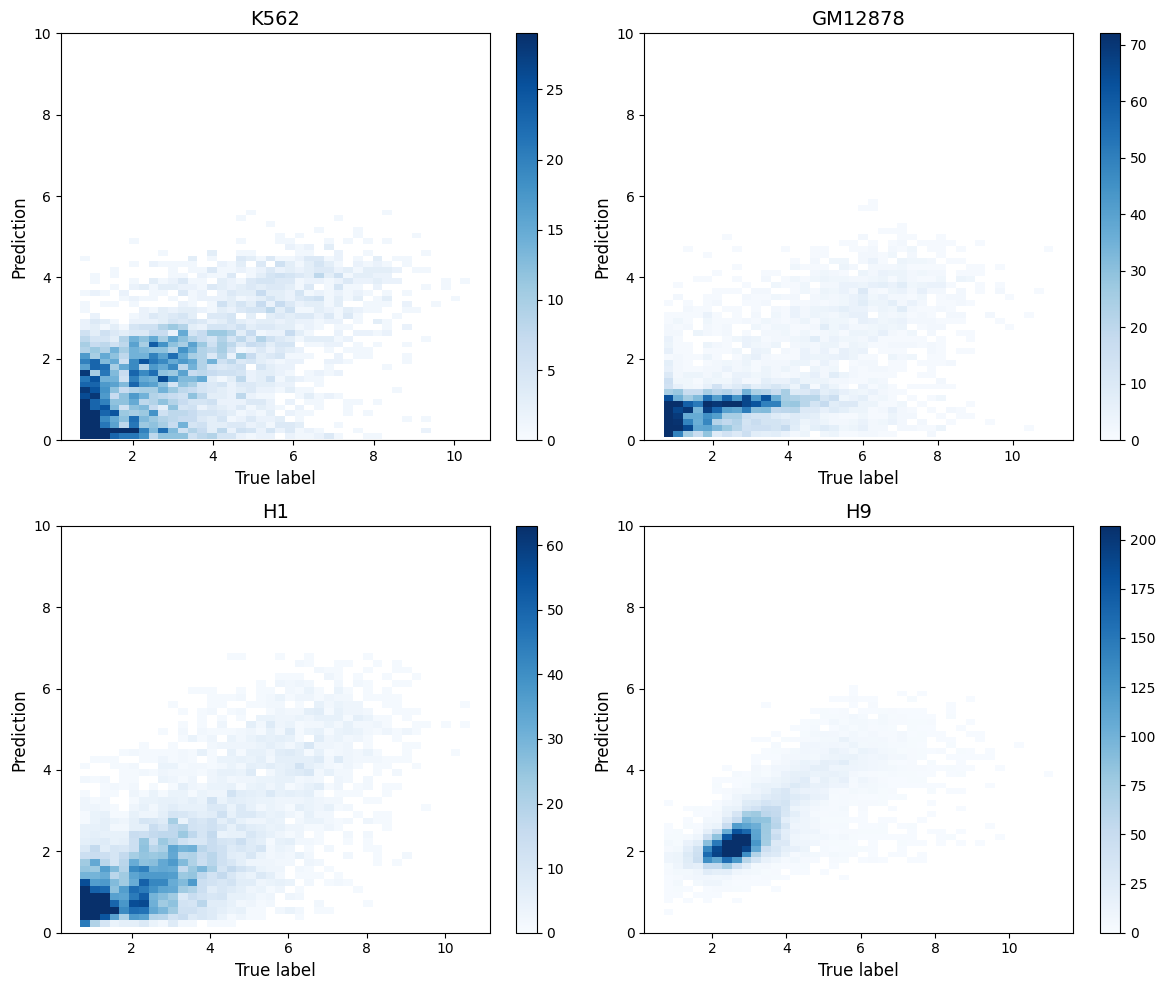

Figure S6. Comparison of model-predicted and experimentally measured CAGE coverage across K562, GM12878, H1, and H9 cell types.

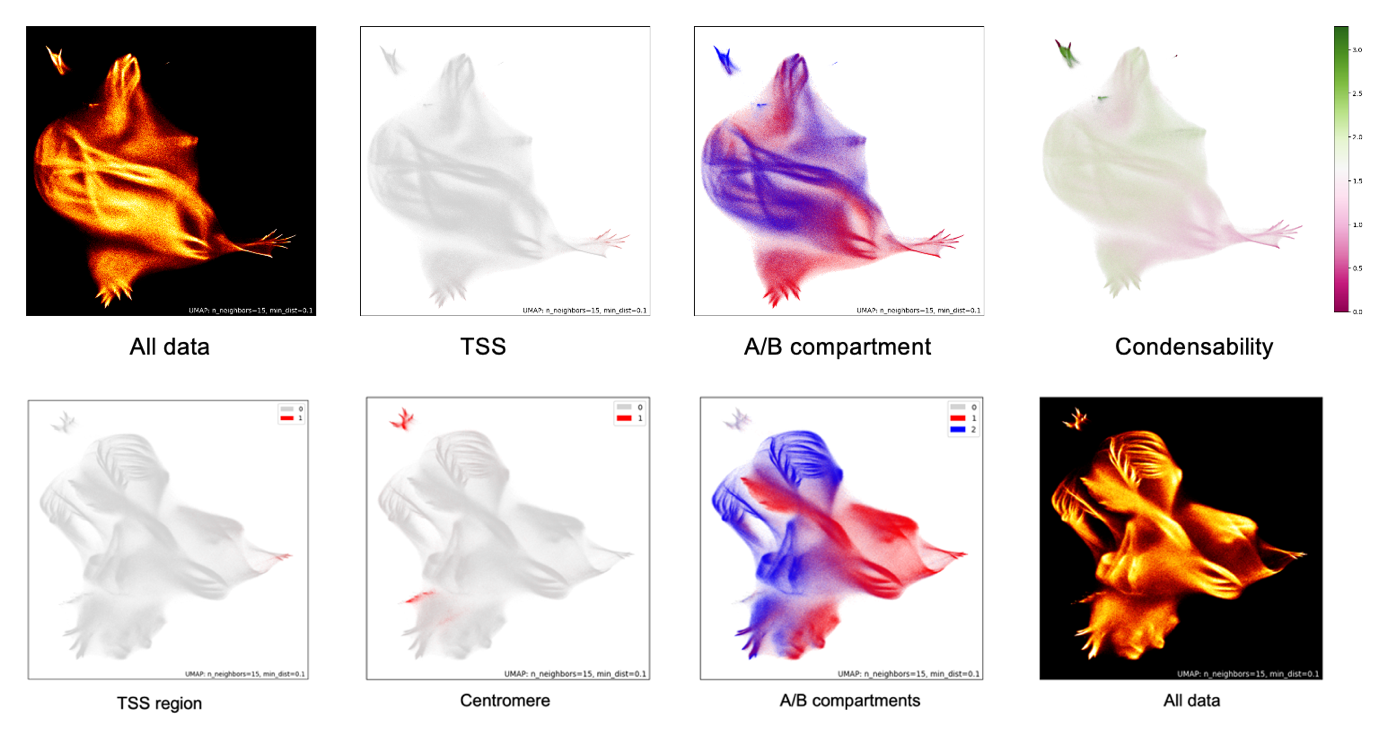

Figure S7. UMAP projection of sequence embedding vectors based on the H1 cell, while each mark represents a 100bp DNA sequence.

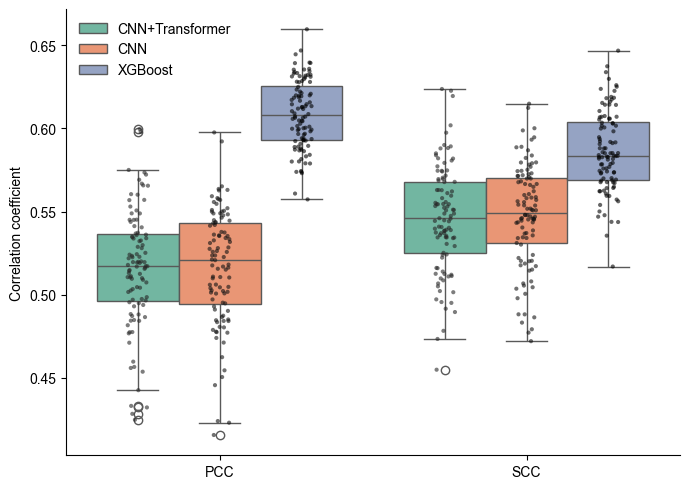

Figure S8. Performance comparison of models based on the PCC and SCC using the GM12878 cell.

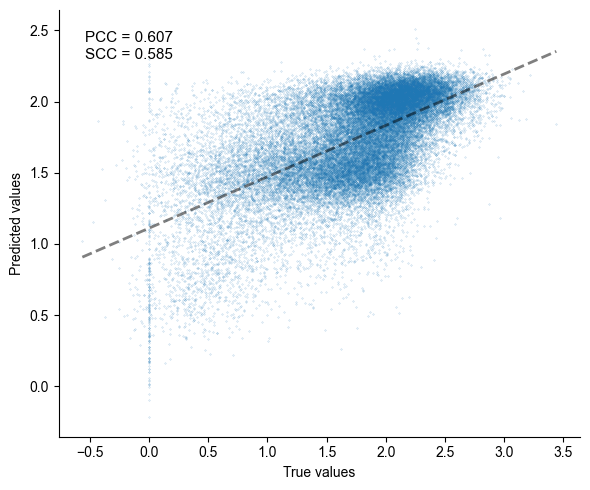

Figure S9. Scatter plot showing the relationship between predicted and true values generated by the XGBoost model for the GM12878 cell line. Predictions were obtained from the test sets of the 10-repeated 10-fold cross-validation. The fitted regression line is shown, and the PCC and SCC are annotated.

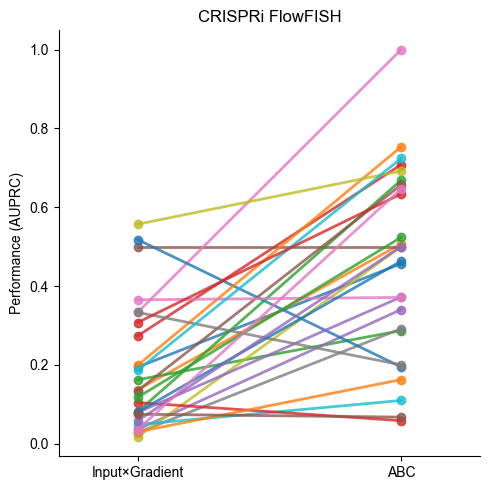

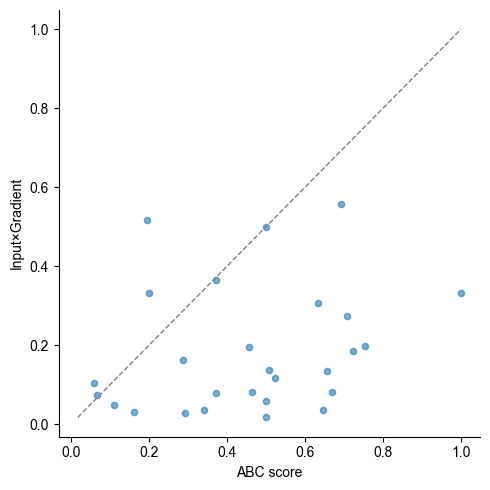

Figure S10. Line plot comparing the performance of enhancer identification using contribution scores versus ABC scores. (b) Dot plot illustrating the AUPRC results for each gene, comparing our model with ABC scores.

Table S1. The accession ID of the training dataset.

| Cell lines | DNase | H3K4me3 | H3K27ac | H3K36me3 | H3K9me3 | H3K4me1 | H3K27me3 | H4K20me1 | CAGE | Hi-C |
| --- | --- | --- | --- | --- | --- | --- | --- | --- | --- | --- |
| K562 | ENCFF972GVB | ENCFF525ZRM | ENCFF381NDD | ENCFF440XMD | ENCFF601JGK | ENCFF761XBZ | ENCFF139KZL | ENCFF605FAF | ENCFF754FAU | 4DNFIR79UBF2 |
| GM12878 | ENCFF960FMM | ENCFF287HAO | ENCFF469WVA | ENCFF831WJT | ENCFF392LMQ | ENCFF564KBE | ENCFF039JOT | ENCFF309OEW | ENCFF002MHE | 4DNFI3PNAYBK |
| H1 | ENCFF573NKX | ENCFF760NUN | ENCFF919FBG | ENCFF370UNF | ENCFF350QBK | ENCFF396RXV | ENCFF380KPI | ENCFF613HTY | ENCFF930AEF | 4DNFICRGVN5L |
| HepG2 | ENCFF842XRQ | ENCFF500VAH | ENCFF022TZG | ENCFF247LOP | ENCFF485BCI | ENCFF576YVM | ENCFF529HLC | ENCFF554JDF | ENCFF933UYY | 4DNFIS6HAUPP |
| H9 | ENCFF405MMH | ENCFF085UOM | ENCFF429TIE | ENCFF728GNA | ENCFF832TVU | ENCFF276TJN | ENCFF577WGF | ENCFF548ENQ | CNhs11917 | 4DNFIT5YVTLO |
| HeLa-S3 | ENCFF194NJI | ENCFF460QIU | ENCFF658XKZ | ENCFF157RUQ | ENCFF658AOG | ENCFF860GGZ | ENCFF614HNF | ENCFF116YMZ | ENCFF530VDY | 4DNFIBM9QCFG |
| IMR-90 | ENCFF971HXR | ENCFF376ZIM | ENCFF699OAR | ENCFF519FHW | ENCFF066MEE | ENCFF221MJG | ENCFF366BVS | ENCFF295EWC | ENCFF615AOF | 4DNFIJTOIGOI |
| HCT116 | ENCFF906MKW | ENCFF964OOU | ENCFF787LMI | ENCFF024LGD | ENCFF254TIW | ENCFF239FXT | ENCFF717ZKL | ENCFF349NQI | E-MTAB-13647 | 4DNFIP8RKGDG |

Table S2. Pearson correlation coefficients of our model across tasks and cell types.

|  | DNase | H3K4me3 | H3K27ac | H3K36me3 | H3K9me3 | H3K4me1 | H3K27me3 | H4K20me1 | CAGE |
| --- | --- | --- | --- | --- | --- | --- | --- | --- | --- |
| K562 | 0.6126 | 0.7662 | 0.5709 | 0.5352 | 0.4211 | 0.6849 | 0.6724 | 0.5469 | 0.6847 |
| GM12878 | 0.3429 | 0.6378 | 0.4395 | 0.558 | 0.2897 | 0.3709 | 0.5044 | 0.3961 | 0.6659 |
| H1 | 0.6255 | 0.8664 | 0.5324 | 0.5308 | 0.4917 | 0.5282 | 0.7539 | 0.4381 | 0.6793 |
| H9 | 0.7305 | 0.8443 | 0.5348 | 0.5072 | 0.4602 | 0.5926 | 0.7613 | 0.4185 | 0.7219 |
| HCT116 | 0.4563 | 0.7893 | 0.4096 | 0.5375 | 0.422 | 0.3766 | 0.4391 | 0.5053 | 0.7869 |
| HepG2 | 0.6679 | 0.7019 | 0.501 | 0.6408 | 0.5949 | 0.4427 | 0.5909 | 0.4104 | 0.6726 |
| HeLa-S3 | 0.2607 | 0.7678 | 0.57 | 0.593 | 0.4379 | 0.3755 | 0.483 | 0.4087 | 0.5805 |
| IMR-90 | 0.2798 | 0.7748 | 0.4221 | 0.6287 | 0.558 | 0.4644 | 0.5091 | 0.5581 | 0.6677 |

Table S3. Spearman correlation coefficients of our model across tasks and cell types.

| Cell lines | DNase | H3K4me3 | H3K27ac | H3K36me3 | H3K9me3 | H3K4me1 | H3K27me3 | H4K20me1 | CAGE |
| --- | --- | --- | --- | --- | --- | --- | --- | --- | --- |
| K562 | 0.7470 | 0.4324 | 0.4261 | 0.5198 | 0.4225 | 0.5082 | 0.6649 | 0.5439 | 0.4790 |
| GM12878 | 0.4698 | 0.4188 | 0.3106 | 0.4375 | 0.3247 | 0.3999 | 0.5450 | 0.4208 | 0.5086 |
| H1 | 0.5491 | 0.4359 | 0.5255 | 0.4101 | 0.4283 | 0.5011 | 0.3555 | 0.4192 | 0.5392 |
| H9 | 0.8791 | 0.4635 | 0.6076 | 0.4140 | 0.4212 | 0.6685 | 0.3855 | 0.4764 | 0.5789 |
| HCT116 | 0.3478 | 0.2708 | 0.4054 | 0.4666 | 0.4332 | 0.4459 | 0.4789 | 0.4867 | 0.4620 |
| HepG2 | 0.6705 | 0.4655 | 0.4035 | 0.5564 | 0.5945 | 0.5304 | 0.5807 | 0.3869 | 0.5249 |
| HeLa-S3 | 0.2443 | 0.5645 | 0.4946 | 0.4742 | 0.4481 | 0.4086 | 0.4937 | 0.4207 | 0.2387 |
| IMR-90 | 0.6159 | 0.3359 | 0.4754 | 0.5793 | 0.5518 | 0.5358 | 0.5475 | 0.6129 | 0.4719 |

Table S4. Performance comparison of machine learning methods for condensability prediction based on the GM12878 cell.

| Methods | PCC | SCC |
| --- | --- | --- |
| XGBoost | 0.6085 (0.0207) | 0.5862 (0.0242) |
| CNN | 0.5166 (0.0348) | 0.5475 (0.03094) |
| CNN+Transformer | 0.5157 (0.0353) | 0.5455 (0.0323) |

*Values are presented as mean (standard deviation) across 10 repeated 10-fold cross-validation.

Table S5. PCC performance of cross-cell line analysis.

| Cell lines | K562 | GM12878 | H1 | H9 | HCT116 | HepG2 | HeLa-S3 | IMR-90 |
| --- | --- | --- | --- | --- | --- | --- | --- | --- |
| K562 | 0.618784 | 0.449081 | 0.453026 | 0.364410 | 0.436521 | 0.481899 | 0.450796 | 0.461453 |
| GM12878 | 0.370483 | 0.489317 | 0.367832 | 0.328588 | 0.362954 | 0.388729 | 0.394371 | 0.371844 |
| H1 | 0.411714 | 0.388578 | 0.613384 | 0.533731 | 0.406392 | 0.424855 | 0.401572 | 0.441264 |
| H9 | 0.344724 | 0.356651 | 0.540544 | 0.630463 | 0.355034 | 0.369773 | 0.368976 | 0.393330 |
| HCT116 | 0.366117 | 0.367894 | 0.392086 | 0.320883 | 0.553863 | 0.442612 | 0.397321 | 0.374902 |
| HepG2 | 0.462372 | 0.442916 | 0.457054 | 0.391675 | 0.483393 | 0.590595 | 0.442020 | 0.467650 |
| HeLa-S3 | 0.404758 | 0.421811 | 0.388042 | 0.340909 | 0.428941 | 0.419205 | 0.506683 | 0.414592 |
| IMR-90 | 0.405958 | 0.395737 | 0.442194 | 0.383764 | 0.382806 | 0.423616 | 0.427816 | 0.554456 |

**Machine learning and deep learning methods for condensability prediction**

**XGBoost**

This model was implemented using the Python version of the XGBoost library (https://xgboost.readthedocs.io/en/stable/index.html). The model consisted of 200 boosting iterations, with each tree constrained to a maximum depth of 6. A learning rate of 0.05 was applied, and stochastic subsampling was employed by sampling 80% of training instances and features per tree. Histogram-based tree growth was adopted to accelerate training. The inputs are the embeddings generated from the 10kb DNA sequences.

**CNN model**

We implemented a convolutional neural network-based model to predict 10 continuous outputs from sequence-derived embeddings. The model input was a generated embedding of shape (1000, 128), where 1000 corresponds to 1,000 genomic bins at a 100-bp resolution, and 128 denotes the embedding dimension. The model first applied three one-dimensional convolutional layers with ReLU activations. Each convolutional block was followed by max-pooling. This produced a fixed-length representation for each input sample. Finally, a fully connected layer was used to map the pooled representation to 10 continuous output values. The model was trained using mean squared error as the loss function for regression. We trained the model for up to 10 epochs with a learning rate of 1e-3 and a batch size of 32.

**CNN_transformer model**

The CNN_transformer model integrates a CNN with a transformer encoder to predict condensability from DNA sequence embeddings. The input to the model is the same as the CNN model. The model first applies a three-layer 1D CNN with ReLU activations, each followed by max-pooling layers of sizes 4, 5, and 5, respectively. The CNN output is then linearly projected into a higher-dimensional space to serve as input to a transformer encoder. The transformer consists of 2 layers, each with 4 attention heads and a feedforward dimension of 1024, enabling the model to capture long-range dependencies within the sequence. Finally, the transformer output is mean-pooled across the sequence dimension and passed through a fully connected layer to generate a 10-dimensional prediction vector corresponding to the target values. This architecture effectively combines local pattern extraction via CNNs with global context modeling through the transformer.
